# Single-cell transcriptional landscape of muscle-derived stem/progenitor cells reveals hallmarks of aging and rejuvenation

**DOI:** 10.64898/2026.04.28.721405

**Authors:** Kavitha Mukund, Seth D. Thompson, Chelsea L. Rugel, Kamil K. Gebis, Richard L. Lieber, Jeffrey N. Savas, Shankar Subramaniam, Mitra Lavasani

## Abstract

Muscle-derived stem/progenitor cells (MDSPCs) are an adult stem cell population with demonstrated regenerative and rejuvenative potential distinct from other muscle progenitor cells. However, their molecular identity and developmental status remain poorly defined. Using single-cell transcriptomics and proteomics, we comprehensively profiled murine MDSPCs across age groups. We show that MDSPCs exist along a transcriptional continuum of maturation—ranging from metabolically active, proliferative early-stage cells to late-stage, lineage-committed myogenic populations. While lacking canonical pluripotency markers, early-stage MDSPCs express gene programs associated with embryonic progenitor identity, suggesting a non-canonical, multipotent-like state. These features distinguish them from both satellite cells and committed myoblasts. Aging reshapes this continuum by reducing stemness-associated signatures while enhancing differentiation programs and oxidative stress. Our identification of distinct MDSPC states provide critical insights into mechanisms that underly tissue regeneration and aging. These findings offer a blueprint for development of future regenerative therapies to combat age-related functional decline.

## Main

Muscle-derived stem/progenitor cells (MDSPCs) have emerged as an important multipotent cell population with significant regenerative potential^1–4^. Isolated using a modified preplate technique based on differential adhesion^5^, MDSPCs are characterized by their capacity for long-term proliferation, self-renewal, and multi-lineage differentiation and repair—including hematopoietic, endothelial, myogenic, osteogenic, chondrogenic, myocardial, and neurogenic lineages^3,6–10^. They have demonstrated therapeutic potential in preclinical models of muscular dystrophy^11^, myocardial infarction^8^, peripheral nerve injury^3^, and aging-related degeneration^1^. They also exhibit resistance to oxidative stress, are immunomodulatory, and can induce neovascularization—all features that contribute to their in vivo neuromusculoskeletal regenerative efficacy^11,1,4^.

Unlike muscle satellite cells, which are relatively well characterized by their expression of *Pax7* and defined myogenic programs^12^, the molecular landscape of MDSPCs remains largely uncharacterized. Here, we comprehensively map the molecular landscape of MDSPCs using single-cell RNA sequencing and protein mass-spectrometry (MS)-based proteomics. Our findings indicate that MDSPCs do not exhibit classical pluripotent signatures but share transcriptional features and regulatory programs with early embryonic progenitors— indicative of a non-canonical, multipotent-like state similar to early developmental stages, offering mechanistic insights into their unique therapeutic potential.

We transcriptionally profiled previously isolated populations of MDSPCs (also known as muscle-derived stem cells, MDSCs), from three young (three-week-old; Y1-Y3) and three old (two-year-old; O1-O3) mice using single-cell RNA sequencing (scRNA-seq) (Fig. 1a, see Methods). The data was processed and integrated using Seurat v5.1 (see Methods), resulting in a final data object containing 20,068 features across 60,226 cells from both young and old samples (Supplementary Fig. 1a, Supplementary Table 1). Clustering analysis identified 18 transcriptionally distinct clusters (clusters 0-17, Fig. 1b), which were further consolidated into eight distinct “groups” (C1-C8, Fig. 1c) based on their unique transcriptional states. Among these, C4 was the largest group (17,659 cells), while C6 was the smallest (1,502 cells) (Fig. 1d). Given limited prior research in characterizing the transcriptional landscape of MDSPCs, we first assessed whether any of the defined groups (C1–C8) exhibited canonical lineage specification programs or stemness-associated transcriptional signatures (Fig. 1e). While groups C4, C5, C7, and C8 showed expression of stemness-associated markers, only C2 expressed muscle lineage markers. No other group showed strong expression of markers associated with any alternative lineage specification (Fig. 1e).

**Figure 1.**
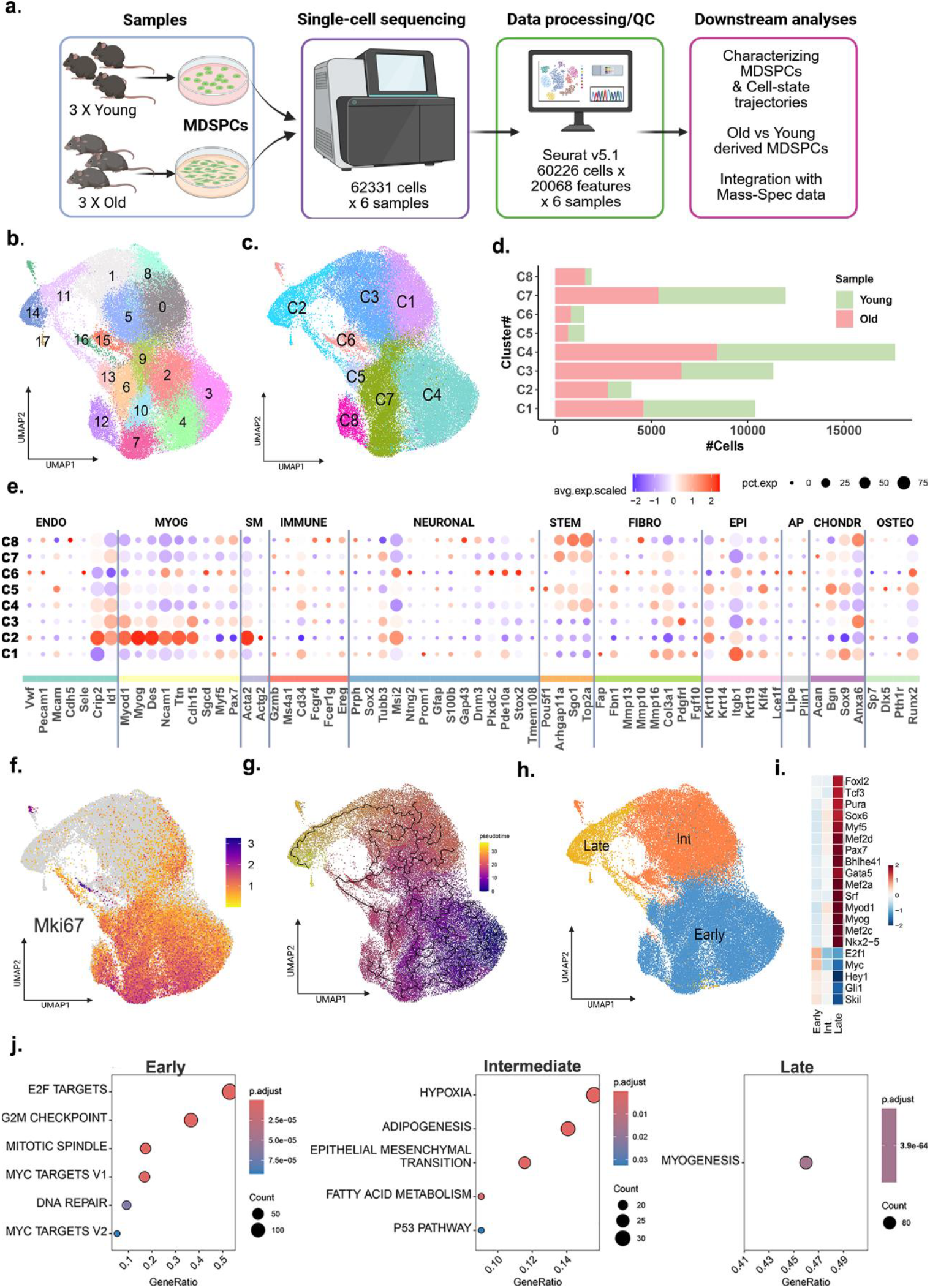
Transcriptional landscape of muscle-derived stem/progenitor cells (MDSPCs). **a**, Schematic overview of the study. **b**, UMAP showing the 18 clusters identified within the integrated UMAP object containing MDSPCs derived from young (21-day-old) and old (2-year-old) mice. **c**, Regrouping of the 18 clusters into 8 distinct groups (C1-C8) based on underlying transcriptional signatures. **d**, Contribution of cells from young and old into groups C1-C8. **e**, A dotplot highlighting the expression of canonical markers across several cellular lineages within groups C1-C8, shows expression of myogenic markers within C2, and stemness within C4/5 and C7/8 **f**, Expression of the proliferation marker Mki67 which indicates cellular proliferation. **g**, Trajectory analysis highlights C2 as being terminal in the pseudotime. **h**, Regrouping of the clusters C1-C8 into three distinct stages–Early, Intermediate (Int), and Late, based on the underlying transcriptional profiles, defined as the maturation axis of MDSPCs. **i**, DecoupleR analysis highlighting the transcription factors driving the expression of genes within cell groups identified in early, intermediate and late stages. **j**. Hallmark enrichment of genes identified as group markers for each of the stages.

To further elucidate the underlying biology of each group, we identified “group markers” that captured the most salient gene expression signatures within each group. Functional enrichment of the larger cell groups, including C4 and C7, showed robust proliferative signatures characterized by elevated *Mki67* expression (Fig. 1f) and enrichment for cell cycle-related pathways, such as mitotic spindle organization, E2F targets, and G2/M checkpoint control. C1 and C3 displayed distinct yet complementary transcriptional programs. Both were enriched for stress-responsive pathways including hypoxia and P53 signaling (Supplementary Fig. 1b), consistent with a transcriptionally plastic state. C3 was further characterized by active metabolic remodeling, with enrichment for fatty acid metabolism, adipogenesis, and mTORC1 signaling alongside myogenesis, suggesting a metabolically primed state. C2 expressed key markers of skeletal muscle maturation, including *Myog* (commitment), *Mymx* (fusion), and *Acta1* (maturation) consistent with a committed myogenic identity^13^ (Supplementary Fig. 1b-c). Trajectory analysis revealed a pseudo-temporal progression across groups, with C4 and C7 being early, and C2 being terminal within this trajectory (Fig. 1g, see Methods). Together, these findings suggest that MDSPCs exist along a continuum of transcriptional states along what we refer to as the “maturation axis”, ranging from proliferative and metabolically active populations to committed myogenic cells.

To decipher the states captured along this maturation axis, we broadly regrouped the eight transcriptional groups (C1-C8) into three stages along this continuum, based on marker expression. These stages represent three transcriptionally distinct states along the maturation axis: Early (C4, C5, C7, and C8), Intermediate (Int) (C1, C3, and C6), and Late (C2) (Fig. 1h). Cells captured within each of these stages are hereafter referred to as Early-, Int-, or Late-MDSPCs (see Methods). To investigate the regulatory mechanisms underlying these three maturation stages, we employed decoupleR^14^ to infer transcription factor (TF) activity. Notably, Late-MDSPCs exhibited increased activity of canonical muscle-specific TFs^15^, including *Myod1, Myog, Myf5, Pax7, Six1* and several *Mefs*. In contrast, Early-MDSPCs were enriched for TFs associated with proliferation, such as *E2F4, Myc, and Ski1* (Fig. 1i). Int-MDSPCs displayed a relatively higher activity for canonical myogenic lineage commitment TFs relative to Early-MDSPCs, but not as high as Late-MDSPCs, consistent with a transitional state. Functional analysis of the maturation stage-specific “group markers” (Supplementary Table 2) supported these findings, as Early-MDSPCs displayed signatures of dynamic cellular reorganization, Int-MDSPCs showed features indicative of plasticity, and Late-MDSPCs showed significant enrichment for myogenesis (Fig. 1j).

Given that these MDSPCs are muscle-derived and exhibit transcriptional features suggestive of cellular plasticity and broad developmental potential along the maturation axis, we next asked whether their transcriptional profiles are recapitulated in muscle stem cells. To address this, we first extracted the top 10% of the group markers identified for each MDSPC maturation stage—yielding 151 for Early, 70 for Int, and 126 for Late (Supplementary Table 3, Supplementary Fig. 2a), and assessed their expression within unipotent muscle stem cells (MuSCs, also known as satellite cells) and committed myoblasts. Transcriptional data for activate and quiescent MuSCs and myoblasts were downloaded from the single-cell muscle atlas (scMuscle, see Methods)^13^. Notably, several of the 126 Late-markers were highly expressed specifically within committed myoblasts, including several troponins, myosin heavy chains, and myogenins (Fig. 2a, Supplementary Fig. 2b). In contrast, unipotent MuSCs exhibited a minimal expression of Early and Int-MDSPC top markers (Fig. 2a), underscoring the distinct transcriptional programs that define the early MDSPC states.

**Figure 2.**
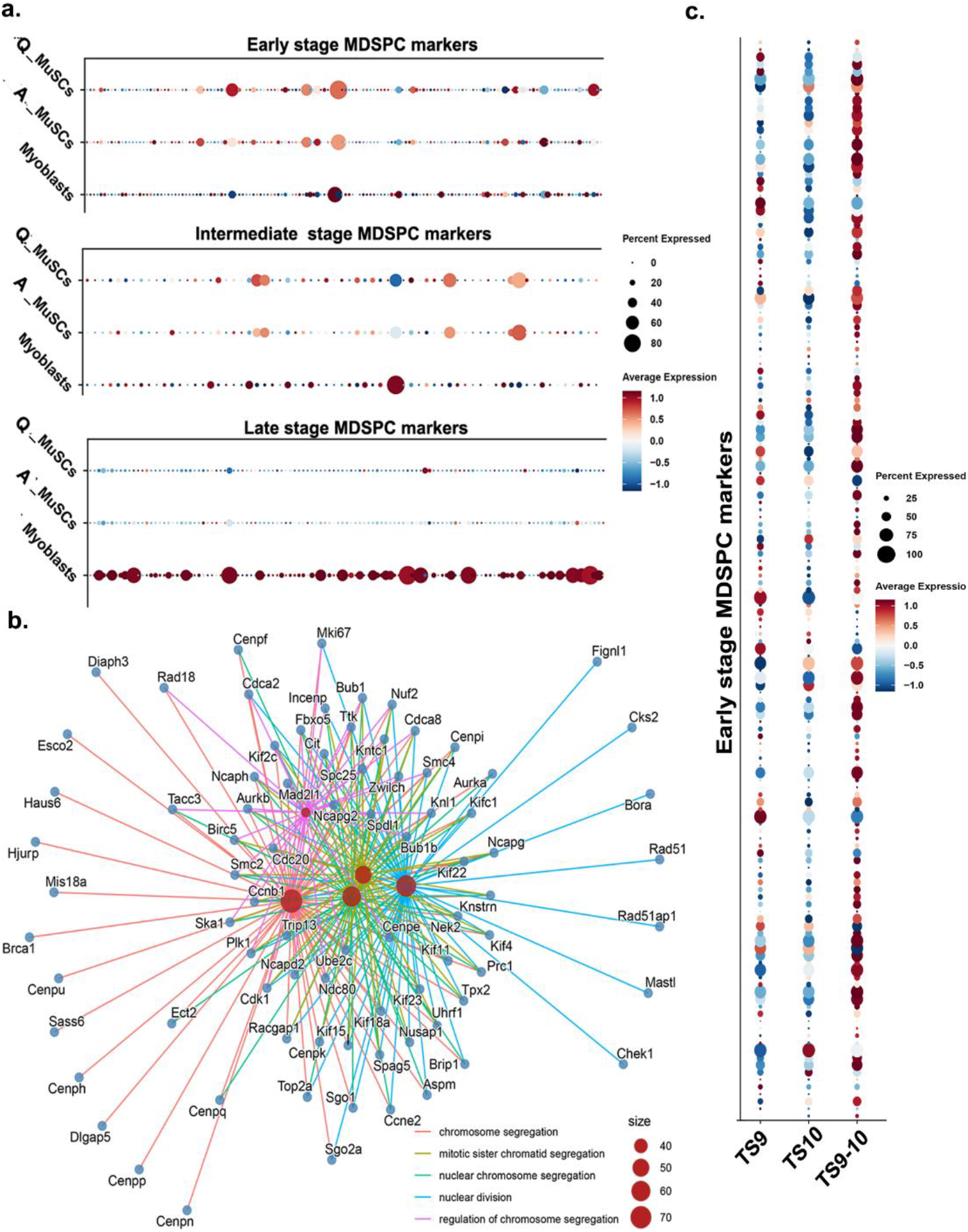
A comparison with muscle stem cells and embryogenesis models. **a**, Expression of Early, Intermediate, and Late-stage markers identified within MDSPCs in quiescent (Q), activated (A) MuSCs (satellite cells) and committed myoblasts extracted from skeletal muscle (see Methods) **b**, A CNET enrichment plot of the Early-stage markers, highlighting the 5 top terms and the genes involved with each term all associated with cellular proliferation and genome maintenance. **c**, Expression of early MDSPC markers in mouse embryogenesis models of gastrulation (pluripotent) at Theiler stages 9 to 10.

Consistent with this, the top markers for Early-MDSPCs (151 genes) displayed an overrepresentation of pathways associated with rapid cell proliferation, lineage priming, and genome maintenance (Fig. 2b). The top markers included genes such as *Birc5, Rad51, Hmgb2, Pclaf, Top2a, Ube2c, Aurkb, Plk1, Cdk1, Bub1, Sgo1, Sgo2a*, and members of *CENP* family, many of which have been shown to play essential roles in genomic stability and cell cycle control in early mouse development^16–18^. The similarity of this Early-MDSPC transcriptional program to proliferative and pluripotent states prompted us to ask whether these features might parallel those observed during early embryogenesis, particularly during the pluripotent stages of mouse development. Utilizing publicly available mouse gastrulation data in pluripotent stages of development, Theiler stages TS9 and TS10 (see Methods), we observed that several Early-MDSPC markers showed significant expression within these stages, and prominently in cells transitioning between TS9 and TS10 (Fig. 2c). This Early-MDSPC to early-gastrulation transcriptional overlap appears to occur despite the absence of canonical pluripotency marker expression within MDSPCs (e.g., *Oct4/Pou5f1, Sox2*, and *Nanog*) (Supplementary Figs. 2c-d). While transcriptional data represent a snapshot in time, further research is warranted to capture and assess the temporal dynamics of canonical pluripotency markers in MDSPCs. These findings, nonetheless, suggest that although MDSPCs are not classically pluripotent, their transcriptional profile mirrors multipotent embryonic progenitors. Collectively, these findings support the existence of a transcriptional continuum among MDSPCs, spanning a range of developmental potentials along the maturation axis, and highlight their unexpected similarity to early embryonic cell states.

Our previous findings have demonstrated that systemic administration of MDSPCs, isolated from young wild-type mice, significantly extends both lifespan and healthspan in progeroid mouse models and ameliorates age-related degeneration in naturally aged mice^1,2,4,19,20^. In parallel, young MDSPCs have been shown to rejuvenate old MDSPCs in co-culture, reversing functional deficits associated with aging^1^. Consistent with these observations, our recent work further identified secreted factors derived from young MDSPCs, compared to old MDSPCs, as key mediators of their pro-angiogenic and immunomodulatory effect^20^, supporting a paracrine mechanism underlying their systemic benefits to naturally aged mice. To gain deeper insights into the transcriptional landscapes distinguishing young from old MDSPCs, we performed differential gene expression analysis between young and old MDSPCs in the context of the aforementioned maturation stages. We note that while both young and old MDSPCs were well represented across the maturation axis, old MDSPCs were disproportionately enriched in the Late MDSPCs (Fig. 3a). This skewed contribution is consistent with a more committed stress-associated transcriptional profile and reduction in stemness, aligning with prior functional data demonstrating proliferative impairment and diminished regenerative efficacy of old MDSPCs in transplantation studies^1^. Together, these findings suggest that age-related shifts along the MDSPC maturation continuum may directly underlie previously observed differences in regenerative capacity and systemic rejuvnation^1^.

**Figure 3.**
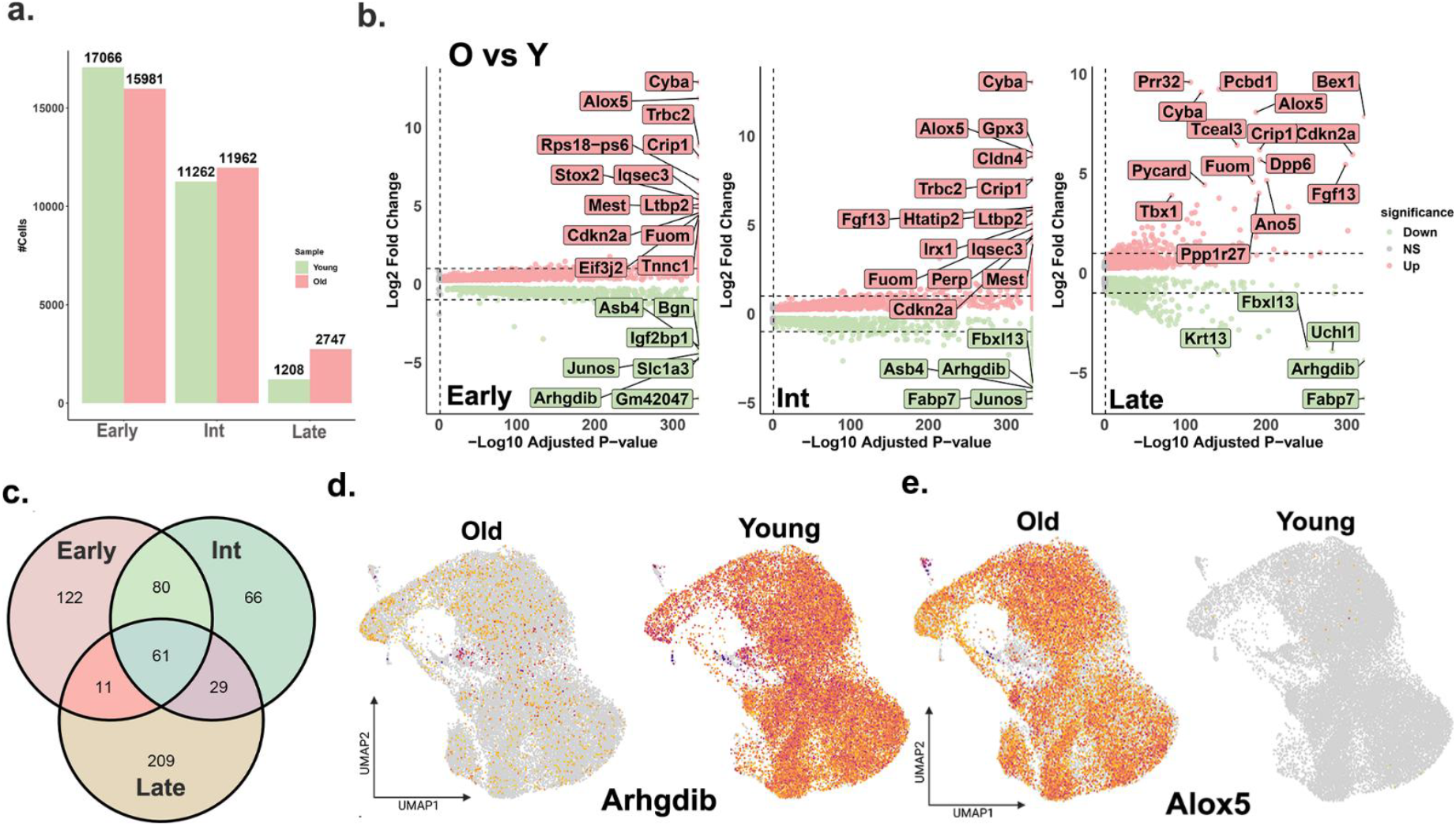
Characterizing the difference between young and old derived MDSPCs. **a**, A barplot summarizing the breakdown of cells derived from young and old mice (*n* = 3 per group), in each of the stages along the MDSPC maturation axis. **b**, Volcano plots highlighting the differential expressed genes (DEGs) identified between old (O) and young (Y) derived MDSPCs in the early-, int- and late-stages along the maturation axis. At a p.adj <0.05, we identified 2432 DEGs, 2333 DEGs and 2102 DEGs in early, int- and late stages respectively. **c**, A Venn highlighting the overlap between DEGs (identified at a revised absolute threshold of log2 fold change >1 (dotted lines in b). **d**, UMAPs highlighting the differential expression in cells for Arhgdib, a gene uniformly albeit, more dominantly expressed within the young, and likewise for **e**, Alox5 is clearly more prominently expressed within old MDSPCs.

Differential gene expression (DEG) of old vs young MDSPCs revealed significant age-related transcriptional changes within each maturation stage, with 2,432 DEGs identified in Early-, 2,333 in Int-, and 2,102 in Late-MDSPCs (Fig. 3b). Broadly, genes upregulated in old MDPSCs were consistently enriched for pathways related to oxidative phosphorylation, p53 signaling, and depending on the maturation stage - myogenic differentiation programs (Supplementary Fig. 3a). In contrast, genes upregulated in young MDSPCs were enriched for pathways associated with epithelial-to-mesenchymal transition (EMT), hypoxic response, and DNA repair (Supplementary Fig. 3b). These contrasting enrichment patterns suggest that aging drives a functional shift in MDSPCs across the maturation axis, with old cells exhibiting stress-response, altered metabolism, and lineage-committed signatures, while young cells retain transcriptional hallmarks of stemness, adaptability, and genomic integrity. This molecular shift may contribute to the age-associated decline in regenerative capacity. Notably, the pathways enriched in young MDSPCs, such as those involved in DNA repair, hypoxic response, and cellular plasticity, may underlie their previously demonstrated ability to extend healthspan and rejuvenate aged MDSPCs.

To emphasize the robust transcriptional differences underlying the observed molecular shifts between young and old, we employed an additional filter to the DEGs, retaining only those with an average log_2_ fold change greater than 1, in addition to the adj p < 0.05. At this revised threshold, we identified 274 DEGs in Early-, 236 in Int-, and 310 in Late-MDSPCs (Fig. 3b, dotted lines, Supplementary Table 4). Of these, 61 DEGs were commonly dysregulated across all maturation stages (Fig. 3c, Supplementary Table 5) and included genes such as *Arhgdib, Cyba, Crip1, Cdkn2a, Alox5, Sema3a/5a, Pdgfc/d, Mysm1, Jun, Fggy, and Fbxl13*. Notably, *Arhgdib* (Fig. 3d) and *Alox5* (Fig. 3e) were among the most significantly dysregulated genes across all maturation stages, differentiating young from old MDSPCs. *Arhgdib*, a Rho GTPase inhibitor known to be abundantly expressed in hematopoietic cells, but relatively less characterized ^21–23^, was highly expressed in young MDSPCs (Fig. 3d); furthermore, young MDSPCs displayed increased expression of genes such as *Sema3a/5a, Pdgfc/d*, and *Mysm1* which are known regulators of vascular development, neural crest migration, and stem cell maintenance^24–28^ across all maturation stages, highlighting conserved pathways shared across mesodermal progenitor populations. In addition to *Alox5*, a key mediator of inflammatory and redox signaling^29^, several other genes associated with oxidative stress, inflammation, and cellular senescence, including *Crip1, Cyba*, and *Mest*, were increased in the old MDSPCs, reflecting progressive loss of homeostatic regulation with age^30–33^.These transcriptional changes were also reflected at the protein level measured by label free MS-based proteomics . Analysis of MS data showed relatively elevated levels of *Alox5* protein in old MDSPCs, whereas proteins such as *Arhgdib, Pdlim5*, and *Igfbp2* were more abundant in young MDSPCs (Supplementary Table 6), supporting coordinated transcriptional–proteomic remodeling across age. Notably, while expression of the canonical myogenic marker *Myod1* was detected in all MDSPCs as expected (Fig. 1e), it was significantly higher in the old (adj p <0.05), particularly within Early- and Int-MDSPCs (Supplementary Table 4). This, taken together with the concurrent expression of the broader transcriptional profile encompassing genes involved in vascular and neural development across all stages, suggests that young MDSPCs possess a more plastic, multipotent identity that extends beyond strict muscle lineage commitment.

Lastly, among the Late-stage DEGs, approximately 68% (209 out of 310) were unique to Late-MDSPCs (Fig. 3c), and significantly increased in the old, including genes such as *Actn2, Actn3, Myom2, Myom3, Myl1, Tcap, Dmd*, and *Cav3*, which encode structural and contractile proteins linked to myofibril assembly consistent with myogenic commitment^15^ (Supplementary Fig. 3c). This, taken together with the cell contributions from old (Fig. 3a), suggests that aging skews MDSPC maturation toward structural specialization at the expense of developmental flexibility.

In total, these data provide the first single-cell transcriptional landscape of MDSPCs defined across a maturation continuum, spanning proliferative, metabolically active early states and transcriptionally committed late myogenic states. This analysis reveals that MDSPCs are not a uniform population but rather a mix of cells in functionally distinct states, with the earliest stages transcriptionally aligned more to multipotent embryonic progenitors rather than canonical muscle stem cells. While we acknowledge that the present study is limited by the use of MDSPCs derived exclusively from female mice, which overlooks potential sex-specific differences in MDSPC biology, and by reliance on cultured rather than freshly isolated cells, such a continuum nonetheless aligns with emerging models of stem cell plasticity, where regenerative potential is tightly linked to metabolic and transcriptional flexibility^34^. Importantly, we also uncover age-associated shifts across this axis: young MDSPCs tend to be enriched for proliferative, hypoxic-response, and DNA-repair programs consistent with a plastic state. While *Arhgdib* emerges as a key regulator, future studies are warranted to gain detailed mechanistic insights into its role in young MDSPCs. In contrast, old MDSPCs upregulate oxidative phosphorylation, p53 signaling, inflammatory mediators (e.g., *Alox5*), with a larger proportion of cells expressing structural genes characteristic of terminal myogenic commitment, reflecting a loss of plasticity and heightened stress- and senescence-associated signatures.

Our findings provide new insights into MDSPCs and their regenerative potential by defining a maturation continuum wherein early states exhibit transcriptional and metabolic programs reflective of high plasticity, while late states are poised for committed myogenesis. The transcriptional heterogeneity described here suggests that MDSPCs are dynamically regulated along axes of stemness, metabolic flexibility, and lineage commitment-with critical implications for regenerative processes in aging tissues. Importantly, these molecular phenotypes correlate with age-associated functional shifts: young MDSPCs display hallmarks of adaptability, genomic integrity and regenerative competence, while old MDSPCs show signatures indicative of cellular stress, metabolic maladaptation, and lineage inflexibility. Together, with our prior evidence that young MDSPCs can preserve tissue homeostasis and counteract age-related degeneration across multiple contexts -- including maintenance of articular cartilage integrity and enhanced tissue repair in models of musculoskeletal injury -- our results suggest that differential MDSPC states may underlie their intrinsic regenerative capacity in aging organisms. These insights offer a roadmap for future efforts to isolate or reprogram specific MDSPC subpopulations with optimal regenerative potential and provide a molecular foundation for therapeutic strategies aimed at mitigating age-related tissue decline across diverse tissues.

## MATERIALS AND METHODS

### Cell isolation and preparation

Young (21-day-old) and old (2-year-old) MDSPCs were previously isolated from the hindlimb skeletal muscle of C57bl/6J mice using a modified preplate technique^5^. In brief, skeletal muscle tissue was enzymatically dissociated with collagenase type-XI (Sigma-Aldrich, C7657), dispase (Invitrogen, 17105-041), and trypsin-EDTA (Invitrogen, 15400-054) to generate a single-cell suspension. Cells were sequentially plated onto collagen type-I (Sigma-Aldrich, C9791) coated flasks, and non-adherent cells were serially transferred to fresh flasks for a total of six preplateing steps over five days to enrich for the slowly adhering MDSPC population. MDSPCs were cultured in collagen type-1 coated flasks containing proliferation media (DMEM [Invitrogen, 11995-073] supplemented with 10% fetal bovine serum [Invitrogen, 10437-028], 10% horse serum [Invitrogen, 26050-088], 1% penicillin/streptomycin [Invitrogen, 15140-122], and 0.05% unfiltered chick embryo extract [Accurate Chemical, MD-004-D-10]) as previously described^5^.

### Library preparation and sequencing

Cell number and viability were analyzed using Nexcelom Cellometer Auto2000 with AOPI fluorescent staining method. Sixteen thousand cells were loaded into the Chromium iX Controller (10X Genomics, PN-1000328) on a Chromium Next GEM Chip G (10X Genomics, PN-1000120) and processed to generate single cell gel beads in the emulsion (GEM) according to the manufacturer’s protocol. The cDNA and library were generated using the Chromium Next GEM Single Cell 3’ Reagent Kits v3.1 (10X Genomics, PN-1000286) and Dual Index Kit TT Set A (10X Genomics, PN-1000215) according to the manufacturer’s manual. Quality control for constructed library was performed by Agilent Bioanalyzer High Sensitivity DNA kit (Agilent Technologies, 5067-4626) and Qubit DNA HS assay kit for qualitative and quantitative analysis, respectively. The multiplexed libraries were pooled and sequenced on Illumina Novaseq 6000 sequencer with 100 cycle kits using the following read length: 28 bp Read1 for cell barcode and UMI, and 90 bp Read2 for transcript.

### Single-cell data processing

Single-cell RNA sequencing was performed at the Northwestern University Sequencing (NUSeq) Core facility, using the Illumina Novaseq 6000 machine. Filtered matrix files for each sample (young Y1-Y3 and old O1-O3) were generated by CellRanger (v7.2.0) at the NUSeq core. All downstream analyses were conducted on these filtered matrices using Seurat v5.1^35^. A total of 62331 cells were captured across six samples. Cells were filtered out if they met any or all of the following criteria-more than 25% mitochondrial content, fewer than 250 expressed genes, transcript count more than 2 standard deviations from the mean, fewer than 500 detected transcripts, or a complexity (log10 genes per UMI) below the 80^th^ percentile. Additionally, genes expressed in fewer than 10 cells or zero counts were excluded. Supplementary Fig 1a shows the number of cells captured per sample, before and after quality control (QC). Post-QC, all samples were merged, log-normalized, and scaled prior to integration. Canonical Correlation Analysis (CCA)-based integration was performed on the merged dataset using 2000 variable features, following the Seurat v5 integration vignette. Fourteen principal components (PCs) were selected (as previously described^36^) and dimensionality reduction was performed UMAP on the first 14 PCs. Cell clustering was carried out using the Louvain algorithm.

### Cellular cluster identity in the integrated dataset

To establish the transcriptional identities of the 18 MDSPC cell clusters, we first identified conserved features for each cluster across all samples within each age group (young and old) using the *FindConservedMarkers* function. We then extracted the top markers by ranking features based on average fold change within age group. To further characterize these clusters, we visualized the expression of the top 20 markers per cluster and performed functional enrichment analysis on the top 500 markers using the enrichR library^37^ (databases: *HDSigDB_Mouse_2021* and *Mouse_Gene_Atlas*) (Supplementary Table 1). Based on the cumulative results of marker expression and enrichment profiles, we grouped the 18 clusters into 8 transcriptionally distinct categories, labeled “Groups C1-C8. “Group markers” were defined as differentially expressed genes (DEGs) that were highly expressed in one group relative to all other groups. This was assessed using the *FindAllMarkers* function in Seurat with Wilcoxon Rank Sum test, applying the following threshold: log fold change = 0.25, expression in at least 25% of cells in the tested group, and adjusted p < 0.05). A similar approach was used to identify “group markers” in early, intermediate, and late stages. The group markers identified for each of the maturation stages were then sorted by average log fold change, and the top 10% were utilized for overlaps with muscle stem cells and mouse organogenesis models.

### Trajectory inference

Trajectories were inferred using reversed graph embedding, available via Monocle 3^38,39^ to model cellular differentiation paths. The preprocessed and reduced CDS object was learned into a graph with learn_graph(), employing default parameters (use_partition = TRUE, close_loop = FALSE) to identify principal graphs without assuming cycles. Root nodes for pseudotime were selected within cluster C4 using order_cells() interactively via graphical interface. Pseudotime values were assigned to each cell along the principal graph, representing progression from the root (pseudotime = 0) to terminal branches.

### Integration of reference data from muscle cell atlas (scMuscle)

We downloaded and utilized the muscle cell atlas (scMuscle) published by McKellar et al^13^, available via Dryad (doi:10.5061/dryad.t4b8gtj34). As our focus was on healthy stem cells from uninjured mice, we subset the dataset for “MuSCs” and “Myoblasts/Progenitors” from samples not exposed to injury agents such as cardiotoxin or notexin. From this, we extracted a total of 33,305 MuSCs including 420 cells annotated as “Activated_MuSCs” and 22,294 as “Quiescent_MuSCs”. Additionally, we retrieved 151 cells annotated as committed myoblasts by the original authors.

### Integration of reference data from mouse organogenesis models

We also incorporated data from a published organogenesis model covering embryonic development stages E6 and E8.5, published by Pijuan-Sala et al^23^ available through the MouseGastrulationData R package^40^. Cells annotated as “doublet” and “stripped” were excluded, resulting in the final dataset of 116,312 high-quality cells spanning 37 distinct cell types, as described in the original study. We extracted cells annotated as Theiler stages 9, 10, and 9-10 from the original data for plotting as they represent the pluripotent stages of gastrulation, while stages 11-12 represent a more lineage committed/tissue restricted stage of development and were excluded.

### Differential gene expression and enrichment analysis

Differential gene expression analysis (DGEA) between time-point groups was performed using the “FindMarker” function within Seurat v5.0, applying the same thresholds for identifying DEGs as described above. Heatmaps of the DEGs, showing fold changes across the relevant conditions, were generated using “pheatmap” library in R/Bioconductor. Gene Ontology (GO) enrichment analysis on the identified DEGs was performed using ClusterProfiler v.4.12.1^41^ or enrichR ^37^. For visualization, the top 5-10 enriched GO categories were selected and presented throughout the manuscript and supplementary materials. ClusterProfiler was also utilized to generate enrichment plots for selected GO terms.

### Transcription factor activity and functional enrichment analysis

Transcription factor (TF)-target activity was assessed using decoupleR^14^ available in R/BioC. TF activity was estimated using the run_decoupleR function with the weighted mean method (run_wmean) based on group markers identified in the early, intermediate, and late cells (3,671 genes x 60,226 cells). The top 25 TFs were selected based on their variability across conditions and were subsequently scaled for downstream analysis. A heatmap of the scaled TF activities was generated to visualize TF dynamics, using a blue-to-red color gradient to represent low to high activity. Hierarchical clustering was performed using the “average” linkage method, and custom color breaks were applied to emphasize variation in TF activity across samples.

### Mass spectrometry sample preparation

Young and old MDSPC protein samples (*n* = 3 biological replicates per group, each analyzed in triplicate) were extracted using a chloroform–methanol precipitation method. Protein pellets were resuspended in 8 M urea (Thermo Fisher Scientific, 29700) prepared in 100 mM ammonium bicarbonate solution (Fluka, 09830). The samples were reduced with 5 mM Tris(2-carboxyethyl)phosphine (TCEP, Sigma-Aldrich, C4706; vortexed for 1 h at room temperature), alkylated in the dark with 10 mM iodoacetamide (Sigma-Aldrich, I1149; 20 min at room temperature), diluted with 100 mM ABC and quenched with 25 mM TCEP. Samples were diluted with 100 mM ammonium bicarbonate solution and digested with Trypsin (1:50 dilution; Promega, V5280) for overnight incubation at 37 °C with intensive agitation. The next day, the reaction was quenched by adding 1% trifluoroacetic acid (Fisher Scientific, O4902–100). Samples were desalted using Peptide Desalting Spin Columns (Thermo Fisher Scientific, 89882). All samples were vacuum centrifuged to dry.

### Tandem mass spectrometry

Three micrograms of each sample were loaded using an autosampler with a Thermo Vanquish Neo UHPLC system onto a PepMap Neo Trap Cartridge (Thermo Fisher Scientific, 174500; diameter, 300 µm; length, 5 mm; particle size, 5 µm; pore size, 100 Å; stationary phase, C18) coupled to a nanoViper analytical column (Thermo Fisher Scientific, 164570; diameter, 0.075 mm; length, 500 mm; particle size, 3 µm; pore size, 100 Å; stationary phase, C18) with a stainless-steel emitter tip assembled on the Nanospray Flex Ion Source with a spray voltage of 2,000 V. An Orbitrap Ascend (Thermo Fisher Scientific) was used to acquire all the MS spectral data. Buffer A contained 99.9% H_2_O and 0.1% formic acid, and buffer B contained 80.0% acetonitrile, 19.9% H_2_O with 0.1% formic acid. The chromatographic run was for 2 h in total with the following profile: 0–8% for 6 min, 8% for 64 min, 24% for 20 min, 36% for 10 min, 55% for 10 min, 95% for 10 min and again 95% for 6 min.

We used the Orbitrap HCD-MS2 method for these experiments. The following parameters were used: ion transfer tube temp = 275 °C; Easy-IC internal mass calibration, default charge state = 2; and cycle time = 3 s. Detector type was set to Orbitrap, with a resolution of 60,000, with wide quad isolation; mass range = normal; scan range = 375–1,500 *m/z*; maximum injection time mode = auto; automatic gain control target = Standard; microscans = 1; S-lens RF level = 60; without source fragmentation; and data type = profile. MIPS was set as ‘on’, included charge states = 2–7 (reject unassigned). Dynamic exclusion enabled with *n* = 1 for 60-s exclusion duration at 10 ppm for high and low with exclude isotopes; isolation mode = quadrupole; isolation window = 1.6; isolation Offset = Off; active type = HCD; collision energy mode = Fixed; HCD collision energy type = normalized; HCD collision energy = 25%; detector type = Orbitrap; Orbitrap resolution = 15,000; mass range = normal; scan range mode = auto; maximum injection time mode = auto; automatic gain control target target = standard; microscans = 1; and data type = centroid.

### Mass spectrometry data analysis and quantification

Protein identification/quantification and analysis were performed with Integrated Proteomics Pipeline - IP2 (Bruker; http://www.integratedproteomics.com/) using ProLuCID, DTASelect2 and Census and Quantitative Analysis. Spectrum raw files were extracted into MS1 and MS2 files using RawConverter (http://fields.scripps.edu/downloads.php/). The tandem mass spectra (raw files from the same sample were searched together) were searched against UniProt mouse (downloaded on 26 October 2020) protein databases and matched to sequences using the ProLuCID/SEQUEST algorithm (ProLuCID version 3.1) with a 50-ppm peptide mass tolerance for precursor ions and 600 ppm for fragment ions. The search space included all fully and half-tryptic peptide candidates within the mass tolerance window with no-miscleavage constraint, assembled and filtered with DTASelect2 through IP2. To estimate protein probabilities and false discovery rates accurately, we used a target/decoy database containing the reversed sequences of all the proteins appended to the target database. Each protein identified was required to have a minimum of one peptide with a minimal length of six amino acid residues. After the peptide/spectrum matches were filtered, we estimated that the protein false discovery rates were ≤1% for each sample analysis. Resulting protein lists include subset proteins to allow for consideration of all possible protein isoforms implicated by at least three given peptides identified from the complex protein mixtures. Then, we used Census and Quantitative Analysis in IP2 for protein quantification (Supplementary Table 7, 24416 protein accessions). Static modification was set to 57.02146 C for carbamidomethylation^42^.

### Mapping proteomic data

Label-free semi-quantitative Normalized Spectral Abundance Factor (NSAF), which helps to correct for variations that can arise during the proteomic workflow^43^, leading to more reliable quantitative results, were averaged for young and old, for each protein accession (Supplementary Table 7). Proteins with a statistically significant difference in relative abundance between young and old groups (p < 0.05, 2307) were retained for downstream analysis. Each protein accession was mapped to gene level (extracted from the UniProt description field-GN tag, Supplementary table 7). Where multiple UniProt accessions mapped to the same gene symbol, the isoform with the highest total ion intensity (summed across young and old) was retained as the representative gene-level measurement. To assess age-dependent changes in protein abundance, the ratio of intensities was log_2_-transformed (log2FC). Positive log_2_ values indicated increased protein abundance in old, while negative values indicate increased abundance in young. The resulting gene-level protein abundance estimates were intersected with transcriptomic data to identify genes exhibiting concordant changes at both the transcript and protein levels (Supplementary Table 6).

## Acknowledgments

We would like to thank Jennifer Wai at the Northwestern University Sequencing Core (NUseq) for her assistance with sequencing.

## Author contributions

K.M., S.D.T., S.S. R.L.L. and M.L. designed the study. K.M. and S.D.T. analyzed all the data, K.M. prepared the figures, and wrote the first draft of the manuscript. S.D.T., C.L.R., and M.L. performed all in vitro experiments. J.N.S. and K.K.G conducted the mass-spectrometry experiments. S.S. and M.L provided overall project guidance and secured funding. All authors have read, revised as necessary, and approved the manuscript.

## Data availability

The mass spectrometry proteomics data have been deposited to the ProteomeXchange Consortium with the following identifier: PXD073464 and on the MassIVE repository with the identifier: MSV000100568. The transcriptomics data have been deposited to GEO with the accession GSE324408.

