## Supplemental Figures 1-3 for "Single-cell transcriptional landscape of muscle-derived stem/progenitor cells reveals hallmarks of aging and rejuvenation"

### Supplementary Figures

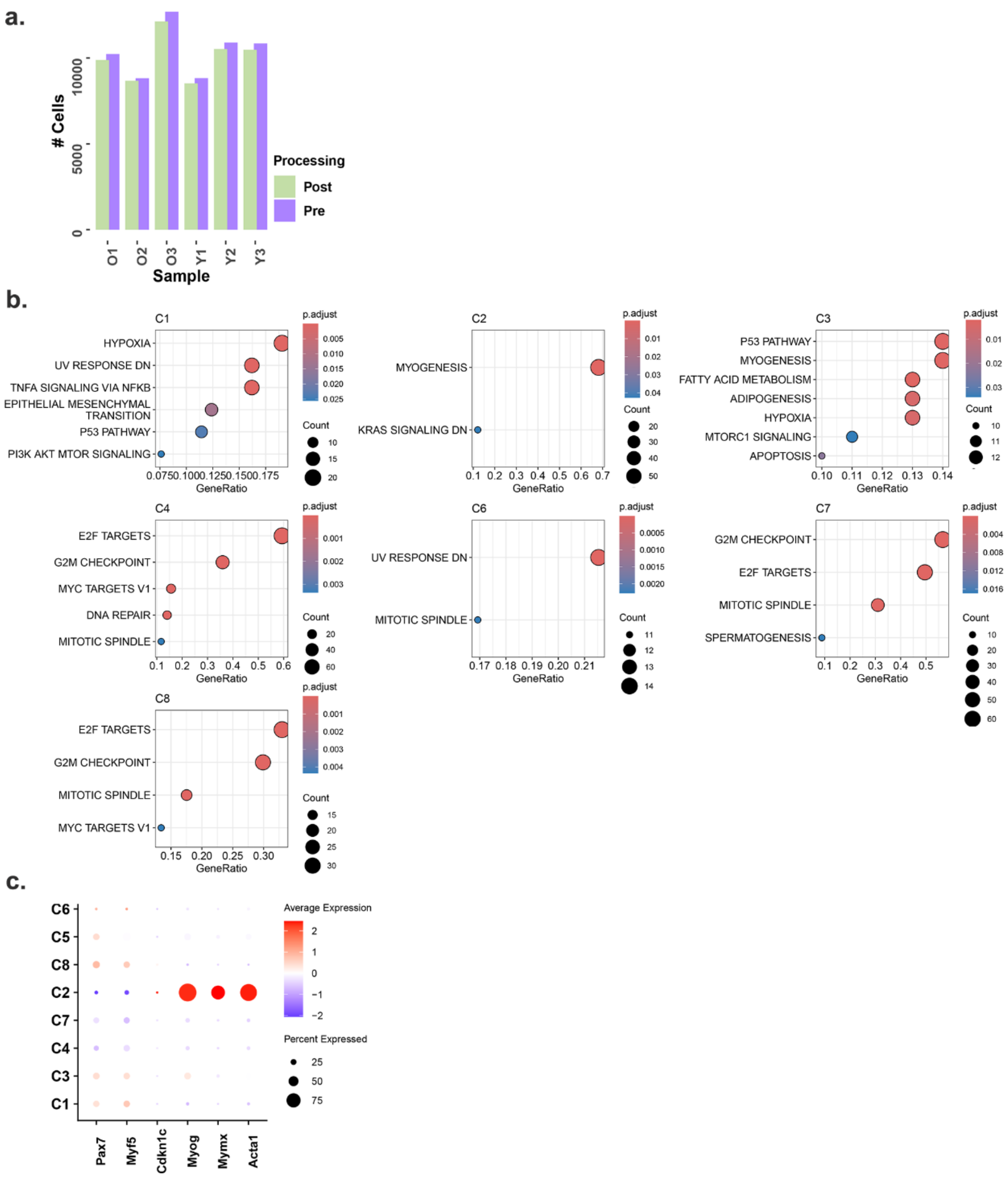


**Supplementary Figure 1 Cell quality control and cluster characterization. a**, Bar plot showing the distribution of cells pre and post quality control (QC) processing. **b**, Hallmark pathway enrichment analysis of group markers identified for clustersC1-C8. **c**, Cluster C2 exhibits expression of skeletal muscle maturation markers, including Myog (commitment), Mymx (fusion), and Acta1 (maturation), consistent with a committed myogenic identity.

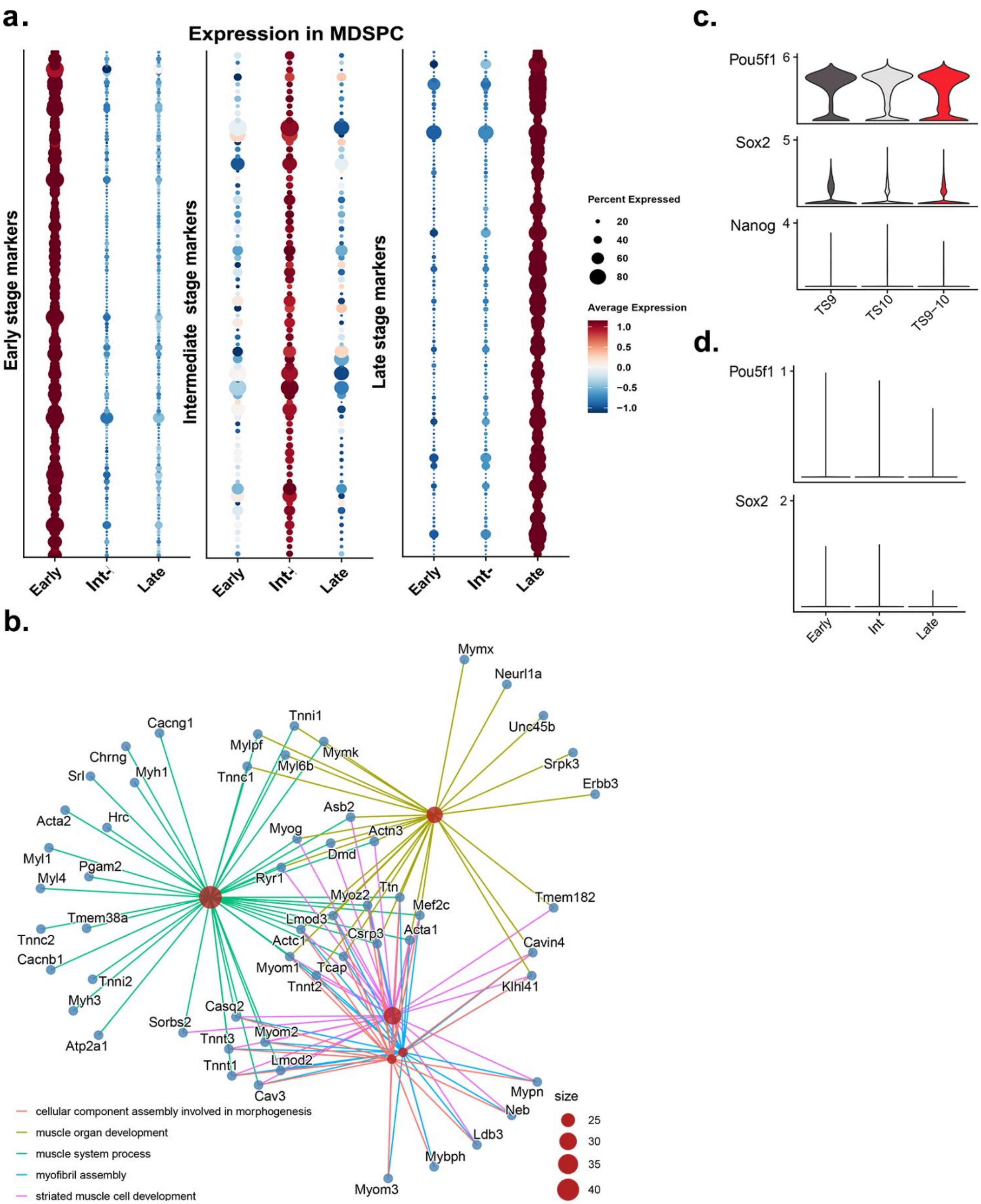


**Supplementary Figure 2 Gene expression and lineage characteristics across differentiation stages**. **a**, A Cnet enrichment plot illustrating biological process and associated genes enriched among late-stage group markers, highlighting commitment to myogenic lineage. **b**, Dotplot showing expression levels of genes identified as group markers for Early, Intermediate and Late stages within our data set. **c**, Expression of canonical pluripotency marker (*Oct4/Pou5f1, Sox2* and *Nanog*) expression with the mouse gastrulation data in Theiler stages TS9-10 and **d**, absence of pluripotency marker expression in MDSPC data across Early, Int, and late stages.

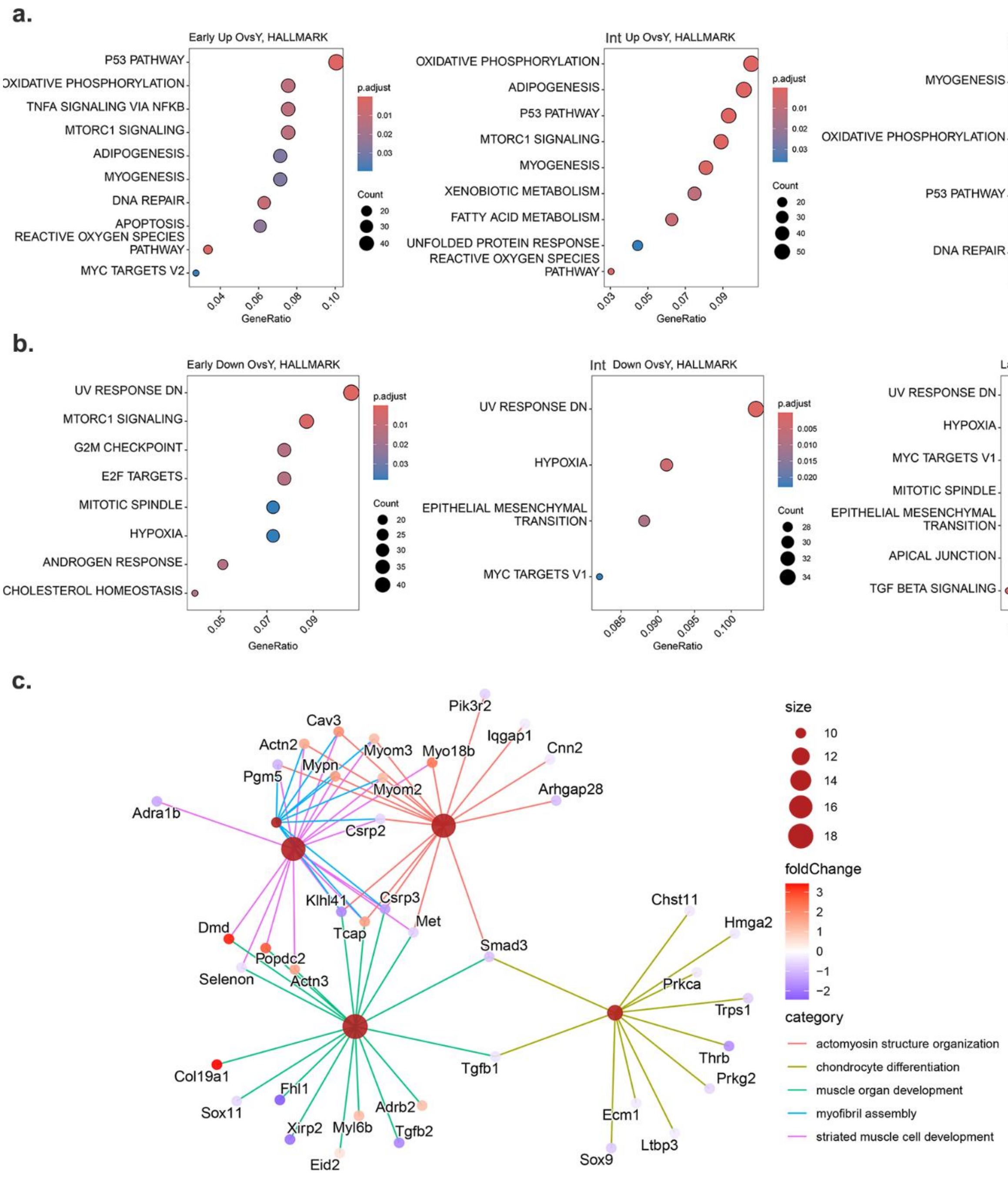


**Supplementary Figure 3 Hallmark pathway enrichment across age-associated gene expression changes. a, b**, Hallmark enrichment analysis of differentially expressed genes (DEGs) that are **a**, upregulated in old and **b**, upregulated in young across Early, Intermediate (Int), and Late stages. **c**, A Cnet enrichment plot for the 209 uniquely expressed DEGs identified exclusively at the late stages of both young- and old-derived MDSPCs.
